# Intranasal Glyphosate-Based Herbicide Administration Alters the Redox Balance and the Cholinergic System in the Mouse Brain

**DOI:** 10.1101/834820

**Authors:** Cristina Eugenia Gallegos, Mariana Bartos, Fernanda Gumilar, Rita Raisman-Vozari, Alejandra Minetti, Carlos Javier Baier

## Abstract

Pesticide exposure is associated with cognitive and psychomotor disorders. Glyphosate-based herbicides (GlyBH) are among the most used agrochemicals, and inhalation of GlyBH sprays may arise from frequent aerial pulverizations. Previously, we described that intranasal (IN) administration of GlyBH in mice decreases locomotor activity, increases anxiety, and impairs recognition memory. Then, the aim of the present study was to investigate the mechanisms involved in GlyBH neurotoxicity after IN administration. Adult male CF-1 mice were exposed to GlyBH IN administration (equivalent to 50 mg/kg/day of Gly acid, 3 days a week, during 4 weeks). Total thiol content and the activity of the enzymes catalase, acetylcholinesterase and transaminases were evaluated in different brain areas. In addition, markers of the cholinergic and the nigrostriatal pathways, as well as of astrocytes were evaluated by fluorescence microscopy in coronal brain sections. The brain areas chosen for analysis were those seen to be affected in our previous study. GlyBH IN administration impaired the redox balance of the brain and modified the activities of enzymes involved in cholinergic and glutamatergic pathways. Moreover, GlyBH treatment decreased the number of cholinergic neurons in the medial septum as well as the expression of the α7-acetylcholine receptor in the hippocampus. Also, the number of astrocytes increased in the anterior olfactory nucleus of the exposed mice. Taken together, these disturbances may contribute to the neurobehavioural impairments reported previously by us after IN GlyBH administration in mice.

## 1. INTRODUCTION

Directly accessible from the environment, the olfactory epithelium (OE) is constantly exposed to huge variety of environmental aggressions. Thus, contaminants such as viruses, prions, agrochemicals, metals, or other potentially harmful agents, may access the central nervous system (CNS) via the intranasal (IN) pathway, causing chronic damage to the brain and leading to neuropathological disorders (Maher et al., 2016; 2012; Rey et al., 2016; Rey et al., 2018).

Pesticides are extensively used throughout the world and abundant evidence has linked pesticide exposure and health impairment, in particular, involving the nervous system. (Kamel and Hoppin, 2004; Le Couteur et al., 1999; Richardson et al., 2014). Glyphosate (Gly) [N-(phosphonomethyl) glycine] is the active ingredient of many broad-spectrum commercial herbicide formulations and constitutes one of the most commonly worldwide used pesticides for weed control. In Argentina, more than 70 % of the cultivable surface is destined to Gly-tolerant corn and soybean (GRAIN, 2009; Trigo, 2011). Considering that these crops are sprayed with 200 million liters of Gly per year (Teubal, 2005; Teubal et al., 2009; Aparicio et al., 2013) it is feasible that the population near to agricultural fields inhale the Gly sprays. The aerial spraying with Gly-BHs was related to dermatological and respiratory illnesses, increased number of miscarriages (Camacho and Mejia, 2017) and DNA alterations in blood samples (Paz-y-Mino et al., 2011; Paz-y-Miño et al., 2007). Moreover, the presence of this herbicide contaminating surface water, soil or food has also been documented (Bohn et al., 2014; Peruzzo et al., 2008; Ronco et al., 2016; Van Stempvoort et al., 2014). Gly has been considered to be safe for non-target organisms. However, recent evidence about the deleterious effects of Gly exposure on human health is beginning to challenge this concept. Clinical reports on human intoxication with commercial formulations of Gly described negative effects in the nervous system, related to Parkinsonian syndrome and alterations of the globus pallidus and substantia nigra (SN) (Barbosa et al., 2001; Wang et al., 2011), as well as anxiety and short-term memory impairments (Nishiyori et al., 2014). Studies in rodents have demonstrated that both, Gly and Gly-based herbicides (GlyBH), produce neurotoxic effects in the CNS. For instance, the exposure of rats and mice to Gly or GlyBH induced impairments in locomotor activity (Ait Bali et al., 2017; Baier et al., 2017; Gallegos et al., 2016; Hernandez-Plata et al., 2015), anxiety behaviour (Ait Bali et al., 2017; Baier et al., 2017; Gallegos et al., 2016) and recognition memory (Baier et al., 2017; Gallegos et al., 2018). Moreover, several neurochemical and oxidative stress markers were altered in the brain of Gly and GlyBH exposed rodents (Ait Bali et al., 2017; Cattani et al., 2017; Cattani et al., 2014; Gallegos et al., 2018; Hernandez-Plata et al., 2015; Martinez et al., 2018)

IN exposure of adult mice to a GlyBH decreases locomotor activity, increases anxiety levels, and impairs recognition memory (Baier et al., 2017). However, the precise mechanism of GlyBH neurotoxicity by the IN pathway remains unclear. Therefore, the purpose of the present study was to dissect the possible mechanisms by which IN GlyBH administration exerts its neuropathological effects. To achieve this objective, adult male CF-1 mice were exposed to repeated IN administration of GlyBH (~2 mg/nostrils/day) three days a week, during four weeks (50 mg/kg/day). Oxidative stress markers and the activity of enzymes such as acetylcholinesterase (AChE) and glutamate (GLU) transaminases were determined in specific brain areas related to the neurobehavioural disorders observed in our previous study (Baier et al., 2017). Additionally, markers of the cholinergic and the dopaminergic pathways, as well as of astrocytes, were analysed through immunofluorescence experiments in coronal brain sections relevant to the behavioural alterations described previously (Baier et al., 2017).

## 2. MATERIALS AND METHODS

### 2.1. Animals

Twenty two adult male CF-1 mice (65 days old), weighing ~40 g, from our own breeding center were used in this study. These were maintained under constant temperature (22 ± 1 °C) and humidity (50–60 %) conditions in a 12 h light-dark cycle, with food (Ganave^®^, Alimentos Pilar S.A., Argentina) and water ad libitum. Both, animal care and handling procedures were in agreement with the standards for the care of laboratory animals as outlined in the NIH Guide for the Care and Use of Laboratory Animals (Garber et al., 2011) and approved by the Institutional Animal Care and Use Committee (CICUAE 089/2016, Departamento de Biología, Bioquímica y Farmacia, Universidad Nacional del Sur, Argentina). Special care was taken as well to minimize the number of mice used.

### 2.2. Materials

The herbicide used in this study is a commercial formulation marketed in Argentina as Glifloglex^®^ from Gleba S.R.L., which contains 48 g of Gly isopropylamine salt per 100 cm^3^ product (equivalent to 35.6 % w/v of Gly acid) (Baier et al., 2017; Gallegos et al., 2018; Gallegos et al., 2016) together with an unspecified mix of inerts and adjuvants. Since the GlyBH utilized in the present study is a commercial formulation, a complete list of inert and adjuvants substances is not provided by the company. The MSDS of Glifloglex^®^ could be found in Supplementary Material 1. In this regard, it is necessary to emphasize that in agricultural practices, Gly formulations are used instead of pure Gly. Then, the goal of this study was to use a commercial GlyBH to which the population is actually exposed to. GlyBH was dissolved in saline at a concentration of 96 mg Gly acid/ml (see section 2.3), after which it was administrated by the IN route. The pH value of saline and GlyBH-saline solution was ~ 5.0 – 5.5 (Baier et al., 2017).

### 2.3. Intranasal administration of a GlyBH solution

Male mice were weighed and randomly assigned to the control group (n = 11), which were administrated with 0.9 % NaCl (saline) solution, or to the GlyBH-treated group (n = 11), respectively. For each experimental group, 6 mice were randomly assigned to behavioural and biochemical determinations, whereas the remaining (n = 5) where destined to the immunofluorescence studies. Mice were adapted to daily handling for one week before starting IN administration. The IN administration of GlyBH was performed as described previously (Baier et al., 2017) (Fig. 1A). Briefly, non-anesthetized CF-1 mice were held by the neck and were laid upside down to limit liquid flow down to the trachea. Animals received the equivalent to ~2 mg of Gly acid/day through the nostrils using a micropipette (10 μl solution/nostril/day; ~1 mg Gly acid/nostril/day; i.e. total/day (both nostrils) ~2 mg Gly acid/day), 3 times per week, during 4 weeks, amounting to 50 mg/kg/day of Gly acid. Control mice were similarly administrated with saline. No adverse contact effects were observed after IN administration. The frequency of administration and GlyBH doses were in the range of what was previously described in the bibliography (see table in Supplementary Material 2).

**Figure 1.**
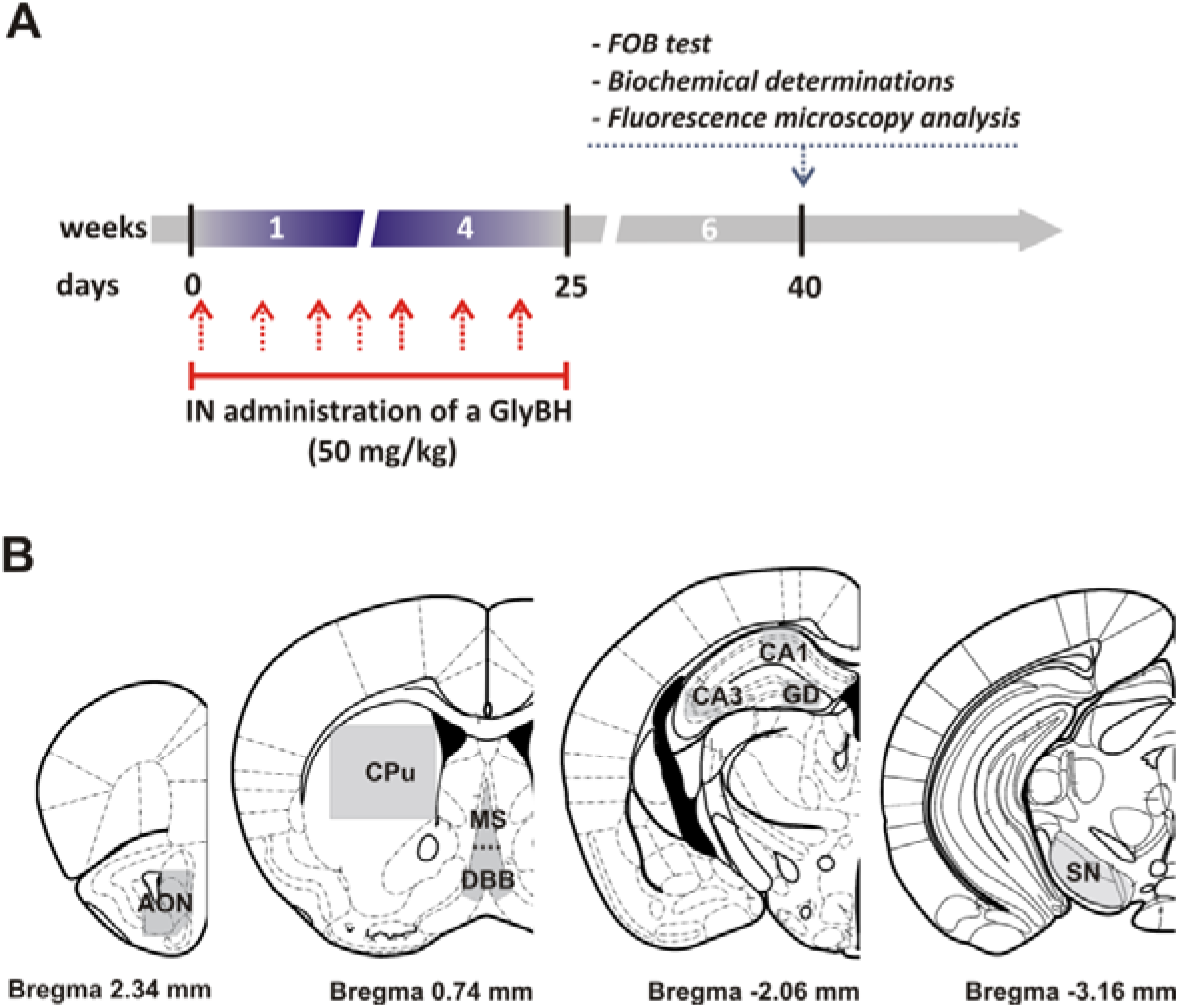
**A)** *Diagram of the experimental design.* **B)** *Representative diagrams of the coronal sections used for immunofluorescence or cell count analysis of mouse brain regions.* Samples included 25 μm sections from: +2.46 mm to +2.22 mm antero-posterior (AP) for AON; +0.98 mm to +0.74 mm AP for MS, DBB and CPu; −1.94 mm to −2.18 AP for HPC; and −3.16 mm to −3.28 for SN AP relative to bregma, respectively, according to Paxinos and Franklin (2001). AON, anterior olfactory nucleus, CPu, striatum; MS, medial septum; DBB, diagonal band of Broca; CA1, CA3 region of the hippocampus (HPC); DG, dentate gyrus of the HPC; SN, substantia nigra. The areas indicated in gray were used for the analyses. The figure is adapted from Paxinos and Franklin (2001).

The systemic no-observed-effect concentration (NOEC) in Sprague– Dawley rats exposed to a GlyBH (Roundup^®^) by inhalation 6 h/day, 5 days/week for 1 month (22 total exposure days) was ? 0.36 mg/L (reviewed in Williams et al. (2000)). Considering that the mouse tidal volume is 4 air L/h (Fairchild, 1972), our GlyBH dose for mice of ~ 40 g weight is about 1.5 (daily) – 3 (accumulated) fold lower than the NOEC.

Additionally, even though the GlyBH dose used in the present work was higher than the GlyBH levels to which the population is normally exposed (Johnson et al., 2005; Solomon, 2016; Williams et al., 2000), it was similar to the doses employed in IN administration studies with other toxins and pesticides (Baier et al., 2017; Prediger et al., 2012; Prediger et al., 2010; Rojo et al., 2007; Rojo et al., 2006; Tristao et al., 2014). Moreover, toxicological studies with pesticides are usually performed at higher doses in order to demonstrate a plausible drug-action mechanism (Baier et al., 2017; Ford et al., 2017).

Body weight, food intake and water consumption were recorded during the IN GlyBH administration time-window.

### 2.4. Functional observational battery (FOB)

The FOB was performed 15 days after the last IN administration (Fig. 1A). FOB included a thorough description of the animals’ appearance, behaviour and functional integrity (US-EPA, 1998). This was assessed through observations in the home cage, but also while animals moved freely in an open field and through manipulative tests. Procedural details and scoring criteria for FOB protocol were performed according to Moser and Ross (1996) and modified for mice (Bras et al., 2010; Ingman et al., 2004; Youssef and Santi, 1997). Briefly, measurements were first carried out in the home cage. The observer recorded each animal’s posture, activity and palpebral closure. The presence or absence of clonic or tonic movements, spontaneous vocalizations and biting were also noted. Then, the animal was removed from its cage, rating the ease of removal and handling. All signs of lacrimation, salivation and piloerection were rated. Other abnormal signs were also recorded. The animal was next placed in an open field arena having a piece of clean absorbent paper on the surface, and allowed to freely explore for 3 min. During that time, the observer ranked for each mouse its arousal, gait score, activity level and rears as well as any abnormal postures, unusual movements, stereotyped behaviours, pelvis elevation and tail position. At the end of the 3 min, the number of fecal boluses and urine pools and presence or absence of diarrhea were recorded. Next, sensorial responses were ranked according to a variety of stimuli (click stimulus using a metal clicker, approach and touch the rump with a blunt object, pinch the tail using forceps, and touch of the corner of the eye and the inside of the ear with a fine object). Also, several motor reflexes were analyzed (flexor, extensor and surface righting reflex). In landing foot splay, the tarsal joint pad of each hind foot was marked with ink and the animal was then dropped from a height of 15 cm onto a recording sheet. This procedure was repeated twice. The distance from center-to-center of the ink marks was measured and the average of the two splay values was used for statistical analysis. Finally, the wire maneuver was carried out. This consisted in suspending the animal from a horizontal wire by its forelimbs and released.

### 2.5. Tissue Preparation

Fifteen days after the last IN GlyBH administration, mice were sacrificed by cervical dislocation (Fig. 1A). Brains were rapidly taken out from the skull and rinsed with ice-cold isotonic saline. The following brain areas were dissected: olfactory bulb (OB), prefrontal cortex (PFC), striatum (CPu), cortex (CTx), hippocampus (HPC) and midbrain (Mb), using the atlas of Paxinos and Franklin (2001) as a guide for tissue dissection. Brain regions were homogenized with disposable homogenization pestles in 10 volumes (1:10, w/v) of ice-cold phosphate buffer saline (PBS; pH 7.4). An aliquot of homogenate was reserved at 4°C for the determination of total thiol content and protein determination, whereas the remaining homogenate was centrifuged and the resultant supernatants were kept at 4°C until determination of enzyme activities. A supernatant aliquot of each sample was reserved for protein determination.

### 2.6. Total thiol content determination

Total thiol (SH) groups were determined using the 5,5’-dithiobis (2-nitrobenzoic acid) method (DTNB). This reagent reacts with SH groups generating a yellow derivative (3-thio-2-nitrobenzoic acid, TNB) with a peak absorbance at 412 nm. Oxidation of free SH groups in proteins leads to the formation of disulfide bonds, which will not react with DTNB. Therefore, the sulfhydryl content is inversely correlated to oxidative damage to proteins (Mazzola et al., 2016). We follow the protocol described by Aksenov and Markesbery (2001) with slight modifications. Briefly, 25 μl of homogenate was added to 1 ml of PBS containing 1 mM EDTA. The reaction was started by addition of 30 μL of DTNB (10 mM in methanol) and incubated for 30 min at room temperature. Samples were then centrifuged, and the absorbance of supernatants was measured at 412 nm. The level of total SH groups were determined from a standard curve prepared with different concentrations of GSH and expressed as μmol TNB/mg protein.

### 2.7. Evaluation of enzyme activities

#### 2.7.1. Catalase (CAT) activity

The activity of the enzyme CAT was determined by the method of Aebi (1984). Reaction was initiated by addition of 0.5 ml H_2_O_2_ (1/10 in PBS) to the reaction mixture containing 100 μl supernatant, 25 μl Triton-X 100 (1/10 in PBS), and 2.4 ml PBS. The decrease in absorbance was recorded for 3 min at 240 nm. The enzyme activity was expressed as the rate constant of a first-order reaction (k) per milligram of protein.

#### 2.7.2. Acetylcholinesterase (AChE) activity

The activity of the enzyme AChE was determined following the Ellman’s method (Ellman et al., 1961). Briefly, an aliquot of each supernatant was incubated with acetylthiocholine iodide (substrate) and 5,5’-dithio-bis(2-nitrobenzoic acid) (DTNB) for 10 min at 30 °C. Enzymatic reaction was stopped by addition of eserine solution and incubation at 0°C for 10 min. The absorbance was measured at 420 nm. AChE activity was calculated from a standard curve prepared with different concentrations of reduced glutathione, and expressed as micromole of thiocholine generated per minute per milligram of protein.

#### 2.7.3. Glutamate oxaloacetate transaminase (GOT) and glutamate pyruvate transaminase (GPT) activities

The activities of GOT and GPT were evaluated by spectrophotometric methods using commercial kits from Wiener Lab. (Rosario, Argentina) following the manufacturers’ indications.

### 2.8. Protein Determination

The protein concentration of total homogenate and the supernatants was measured using the method of Bradford (1976). Bovine serum albumin was used as a standard.

### 2.9. Tissue preparation for immunofluorescence

Fifteen days after the last IN administration of GlyBH (Fig. 1A), mice were deeply anesthetized with a Ketamine/Xylazine mixture (50 and 8 mg/Kg; intraperitoneal injection) and perfused through the cardiac left ventricle, initially with 15 mL of cold physiological saline (0.9 % NaCl solution containing 0.05 % w/v NaNO_2_ plus 50 I.U. of heparin), followed by perfusion with 50 mL of fixative (4 % paraformaldehyde in 100 mM phosphate buffer, PBS, pH 7.4). Brains were then removed, post-fixed for 2 h in 4 % paraformaldehyde, cryoprotected by subsequent overnight immersion in 15 % and 30 % sucrose, and finally stored at −80°C until processing for immunofluorescence. Series of 25-μm-thick coronal sections were cut on a Leica cryostat (Leica, Wetzlar, Germany) as in (Baier et al., 2014; Baier et al., 2015; Debeir et al., 2005). Slices were stored at −20°C in PBS pH 7.4, with 50 % w/v glycerol added as cryoprotector until their use in immunofluorescence studies.

### 2.10. Fluorescence microscopy

Twenty five-μm thick brain sections from both, control and GlyBH treated mice, were selected according to anatomical landmarks corresponding to the (Paxinos and Franklin, 2001) mouse brain atlas (Fig. 1B). To avoid differences in labeling procedure, all sections from control and GlyBH treated mice were simultaneously processed in the free-floating state. In order to block nonspecific binding sites, brain sections were incubated for 1 h at room temperature with 10 % v/v normal goat serum in PBS containing 0.15 % Triton X-100 (PBST). Sections were incubated for 48 h at 4°C with primary antibodies to goat anti-choline acetyltransferase (anti-ChAT, 1/1000, Millipore), rabbit anti-tyrosine hydroxylase (anti-TH, 1/1000, US Biologicals, Salem, MA, USA), rabbit anti-glial fibrillary acidic protein (anti-GFAP, 1/2000, Dako). After five rinses in PBST, sections were incubated for 24 h at 4°C with Alexa Fluor^488^ chicken anti-goat IgG (1/1000, Molecular Probes) or Alexa Fluor^488^ goat anti-rabbit IgG (1/1000, Molecular Probes, Invitrogen, Carlsbad, CA, USA). Additionally, α-Bu ngarotoxi n-Alexa^647^ (αBTX-A^647^, 1/500, Molecular Probes), a toxin that selectively binds to nicotinic acetylcholine receptor (nAChR), mainly to the α7-nAChR subtype (Orr-Urtreger et al., 1997) was used. After further washing in PBST, sections were incubated for 20 min with 0.035 mg/ml Hoechst Stain solution (Sigma). Sections were then washed five times in PBST and finally, mounted on gelatine-coated slides, air dried and coverslipped in 90 % glycerol in PBS until observation.

### 2.11. Image analysis

In order to ensure objectivity, immunolabelled brain section analyses were carried out on coded slides, under blind conditions, with the same standardized observation schedule. Briefly, a single series of sections from each mouse of each group was used for quantification. The brain areas of interest were identified according Paxinos and Franklin (2001) (Fig. 1B), and the boundaries of the target structures in the coronal plane were determined microscopically. Fluorescence images were analyzed using the ImageJ software (National Institutes of Health, Bethesda, MD, USA). For each brain area, fluorescence intensity was measured in the selected field (see below) and the nonspecific background, measured in a region devoid of label, was subtracted. Immunoreactive neurons (ChAT-, αBTX-, TH-or GFAP-positive (+) cells) were counted using a computerized image analysis system. Images were captured from slices using a Nikon Eclipse E-600 microscope (Nikon, Melville, NY, USA) with a K2E Apogee CCD camera driven by CCDOPS software (Santa Barbara Instrument Group, Santa Barbara, CA, USA). In order to quantify ChAT, αBTX, TH or GFAP (+) cells, we used a method similar to that of Traissard et al. (2007). The analysis was made on sections corresponding to +2.46 mm to +2.22 mm for the anterior olfactory nucleus (AON); +0.98 mm to +0.74 mm for the medial septum (MS) and the diagonal band of Broca (DBB), and the CPu; −1.94 mm to −2.18 for HPC; and −3.16 mm to −3.28 for substancia nigra (SN), (AP) relative to the bregma, respectively (Paxinos and Franklin, 2001). For the analysis the relevant area was delimited according to (Paxinos and Franklin, 2001) (Fig. 1B) in each section, whilst the number of neurons obtained was normalized to a fixed area (0.5 mm^2^). Two brain slides from each mouse and brain area were used for the analysis. The section thickness was 25 μm, and the measurements were made on every sixth or tenth serial section (i.e., separated by 150 to 250 μm), to make sure that a same neuron would not be counted twice (Baier et al., 2014; Baier et al., 2015; Debeir et al., 2005).

### 2.12. Statistical analysis

Behavioural test measures in FOB were continuous (providing interval data), ranked (ranks based on a defined scale), descriptive or binary (presence or absence of a sign). These tests were statistically analyzed as follow: continuous data with the Student’s t test, ranked data by means of the Kruskal– Wallis nonparametric test, followed by Mann-Whitney U tests, while descriptive and binary data, were compared using a chi-square test. Biochemical determinations and image analyses were compared using the Student’s t test. Results were expressed as the mean ± SD. A value of p < 0.05 was considered statistically significant. Statistical analyses were carried out using SPSS Statistics, GraphPad Prism and Origin softwares.

## 3. RESULTS

### 3.1. IN GlyBH administration did not alter water and food intake, nor FOB parameters

There were no statistical differences in the consumption of water and food between control and GlyBH exposed groups (Supplementary Fig. S1A and B). In addition, compared to the control group, mice body weight was not affected by IN GlyBH administration (Supplementary Fig. S1C).

Regarding the parameters evaluated by the FOB, repeated subacute IN administration of GlyBH did not produce alterations in home cage or hand-held behaviour, nor in the open field arena observations or during the manipulative tests (Table 1).

**Table 1.** Parameters evaluated in the functional observational battery (FOB) after IN GlyBH administration.

| Endpoints | Control | GlyBH |
| --- | --- | --- |
| **Homecage observations** |  |  |
| Normal body posture (D) | 100 | 100 |
| Activity (R) | 2 | 2 |
| Palpebral closure (R) | 1 | 1 |
| Clonic movements (D) | 0 | 0 |
| Tonic movements (D) | 0 | 0 |
| Biting (D) | 0 | 0 |
| Vocalizations (B) | 0 | 0 |
| **Hand-held observations** |  |  |
| Ease of removal from cage (R) | 1 | 1 |
| Ease of handling (R) | 1 | 1 |
| Salivation (R) | 1 | 1 |
| Lacrimation (R) | 1 | 1 |
| Piloerection (B) | 0 | 0 |
| Normal fur appearance (D) | 100 | 100 |
| Normal respiration (D) | 100 | 100 |
| Normal limb tone (D) | 100 | 100 |
| Normal abdominal tone (D) | 100 | 100 |
| Limb grasping (B) | 100 | 100 |
| **Open field observations** |  |  |
| Activity level (R) | 2,58 | 2,58 |
| Rearing (R) | 1,00 | 1,33 |
| Arousal (R) | 2,75 | 2,50 |
| Normal gait (D) | 100 | 100 |
| Stereotyped behaviours (D) | 0 | 0 |
| Pelvic elevation (R) | 1,92 | 1,83 |
| Normal tail position (D) | 100 | 75 |
| Fecal boluses (C) | 2,17±0,83 | 3,83±0,60 |
| Urine pools (C) | 1,00±0,52 | 0,67±0,49 |
| Diarrhea (B) | 0 | 0 |
| **Manipulative tests** |  |  |
| Approach response (R) | 2,00 | 2,00 |
| Touch response (R) | 2,00 | 2,00 |
| Click response (R) | 2,00 | 2,00 |
| Tail pinch response (R) | 2,00 | 2,25 |
| Palpebral reflex (B) | 100 | 100 |
| Pinna reflex (B) | 100 | 100 |
| Flexor reflex (B) | 100 | 100 |
| Extensor reflex (B) | 100 | 100 |
| Righting reflex (R) | 1 | 1 |
| Landing foot splay (C) | 3,27±0,21 | 3,39±0,15 |
| Wire maneuver (R) | 1 | 1 |

Descriptive (D) and binary (B) data are expressed as percentage of incidence (chi-square test); Ranked (R) data expressed as the mean score of the scale used (Kruskal–Wallis test followed by Mann-Whitney U tests); Continuous (C) data expressed as mean value ± SEM (student's t-test). n = 6 per group.

### 3.2. Repeated IN GlyBH administration alters oxidative stress markers in different mouse brain areas

Alteration of biochemical markers of oxidative damage was observed in specific brain areas of mice exposed intranasally to GlyBH (Fig. 2). Indeed, evaluation of the total thiol content showed a significant decrease in this parameter in OB, CPu, CTx and HPC of IN GlyBH administered animals in comparison with the control group (Figure 3A; p < 0.05). As for the CAT antioxidant system, GlyBH treated mice showed a significant decrease in enzyme activity in several brain territories (PFC, p < 0.005; HPC, p < 0.05; and Mb, p < 0.01) compared to control animals (Fig. 2B).

**Figure 2.**
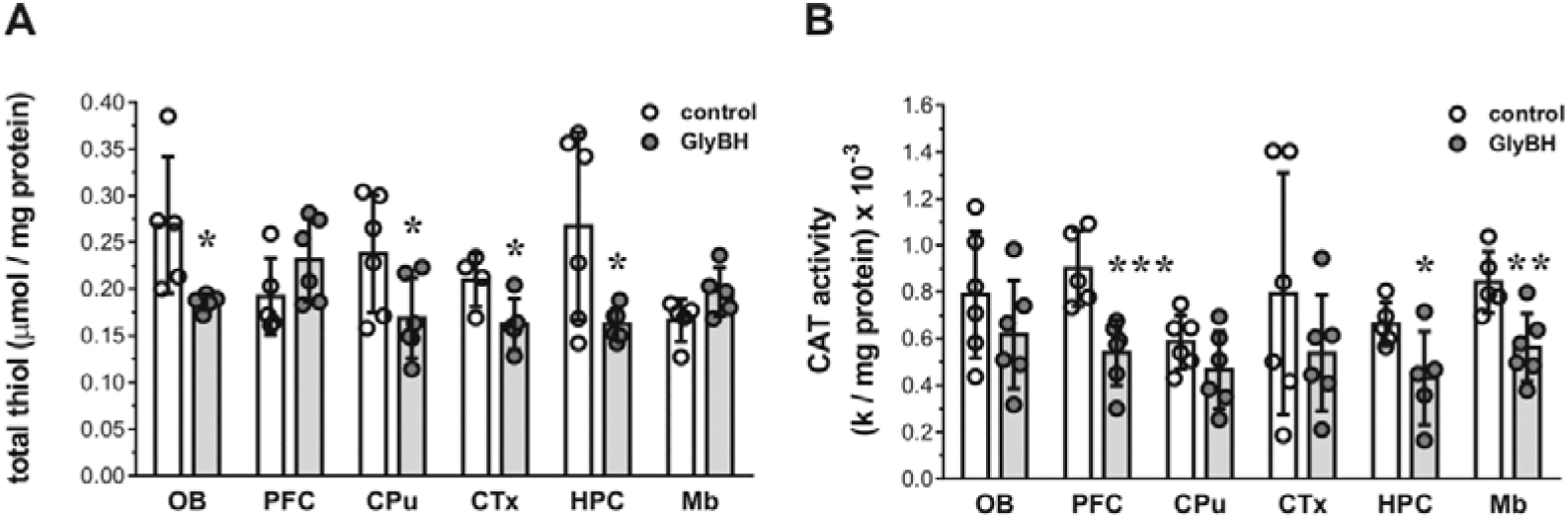
Oxidative stress markers in different brain areas of control and GlyBH treated mice. **A)** Total thiol content at the indicated brain regions from control and GlyBH treated mice. **B)** CAT activity at the indicated brain regions from control and GlyBH treated mice. Bar graphs show mean ± SD, with data points. n = 6 per group. * *p* < 0.05; ** *p* < 0.01; *** *p* < 0.005 compared to the control group (Student’s t test).

### 3.3. Exposure to GlyBH by the IN pathway decreased ChAT expression in the medial septum

Cholinergic neurons in the MS and the DBB are implicated in recognition memory (Okada et al., 2015) and anxiety (Carpenter et al., 2017). Therefore, in order to evaluate whether the cholinergic system in these areas was affected in GlyBH treated mice we performed anti-ChAT immunostaining. As shown in Fig. 3A-B, cell fluorescent intensity (IF, related to protein expression) decreased in the MS of GlyBH exposed group, compared to untreated mice (p < 0.05). Moreover, the number of ChAT (+) cells was lower in herbicide treated mice (p < 0.05) (Fig. 3A and 3C). No alterations were observed in the DBB (Fig. 3B-C). On the other hand, a control area which is not a part of the septo-hippocampal pathway, such as the CPu, did not show changes (Fig. 3B-C).

### 3.4. IN GlyBH administration affected AChE activity in the olfactory bulb

The activity of AChE was studied in the OB, PFC, CPu, CTx, HPC and Mb of control and IN GlyBH exposed mice. Under our experimental conditions, GlyBH administration induced a significant reduction in the activity of this enzyme in the OB (p < 0.05), leaving unaffected the other evaluated brain regions (fig. 3D).

**Figure 3:**
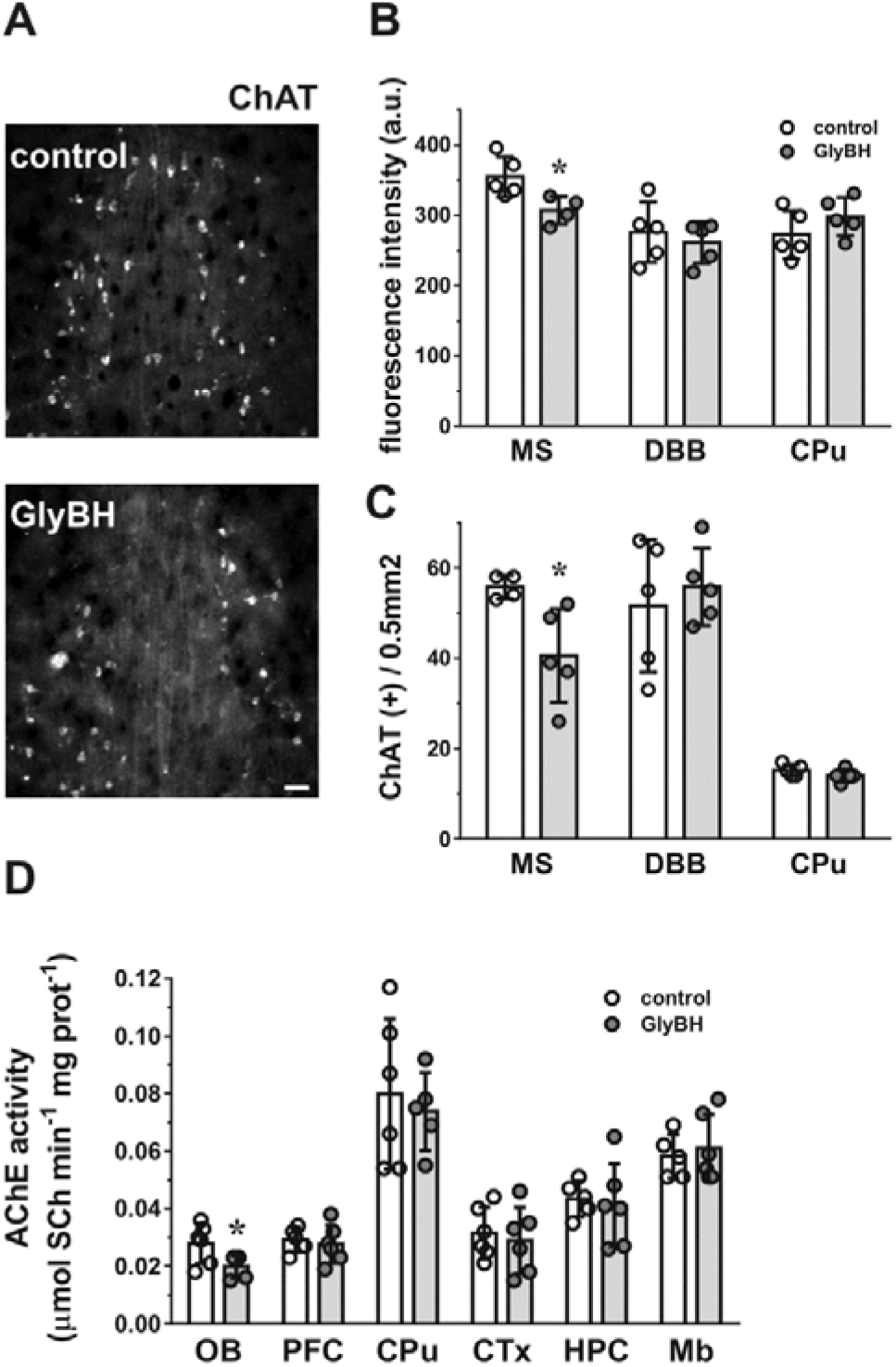
ChAT immunolabelling in the MS, DBB and CPu of control and GlyBH treated mice. **A)** Representative photomicrographs of coronal sections showing ChAT-positive cells in the MS region from control and GlyBH treated mice. Bar, 50 μm. **B)** ChAT fluorescence intensities are expressed as arbitrary units (a.u.). **C)** The number of ChAT-expressing cells was normalized to a fixed area (0.5 mm^2^). Bar graphs show mean ± SD, with data points. * denotes *p* < 0.05, statistically significant different from control mice. Student’s t test, n = 5 mice per group. **D)** AChE activity in the indicated brain regions from control and GlyBH treated mice. Values are mean ± SEM. n = 6 per group. * *p* < 0.05 compared to the control group (Student’s t test).

### 3.5. Intranasally GlyBH exposed mice showed decreased expression of α7-nAChR in the hippocampus

The septo-hippocampal pathway is composed of discrete cholinergic and GABAergic fiber bundles (Khakpai et al., 2013). Modulation of this route is related to anxiety or the detection of novelty (Carpenter et al., 2017; File et al., 1998; Picciotto et al., 2015). In order to evaluate if the cholinergic septo-hippocampal pathway was affected in GlyBH treated mice, the hippocampal expression of α7-nAChR was estimated using fluorescent αBTX. Measurements in the CA1, CA3 and dentate gyrus (DG) hippocampal regions showed that IN administration of GlyBH results in a significant decrease (p < 0.05) in the FI of the αBTX-labelled areas (Fig. 4A-B) without modifying the number of α7-nAChR-expressing cells (Fig. 4A and 4C).

**Figure 4.**
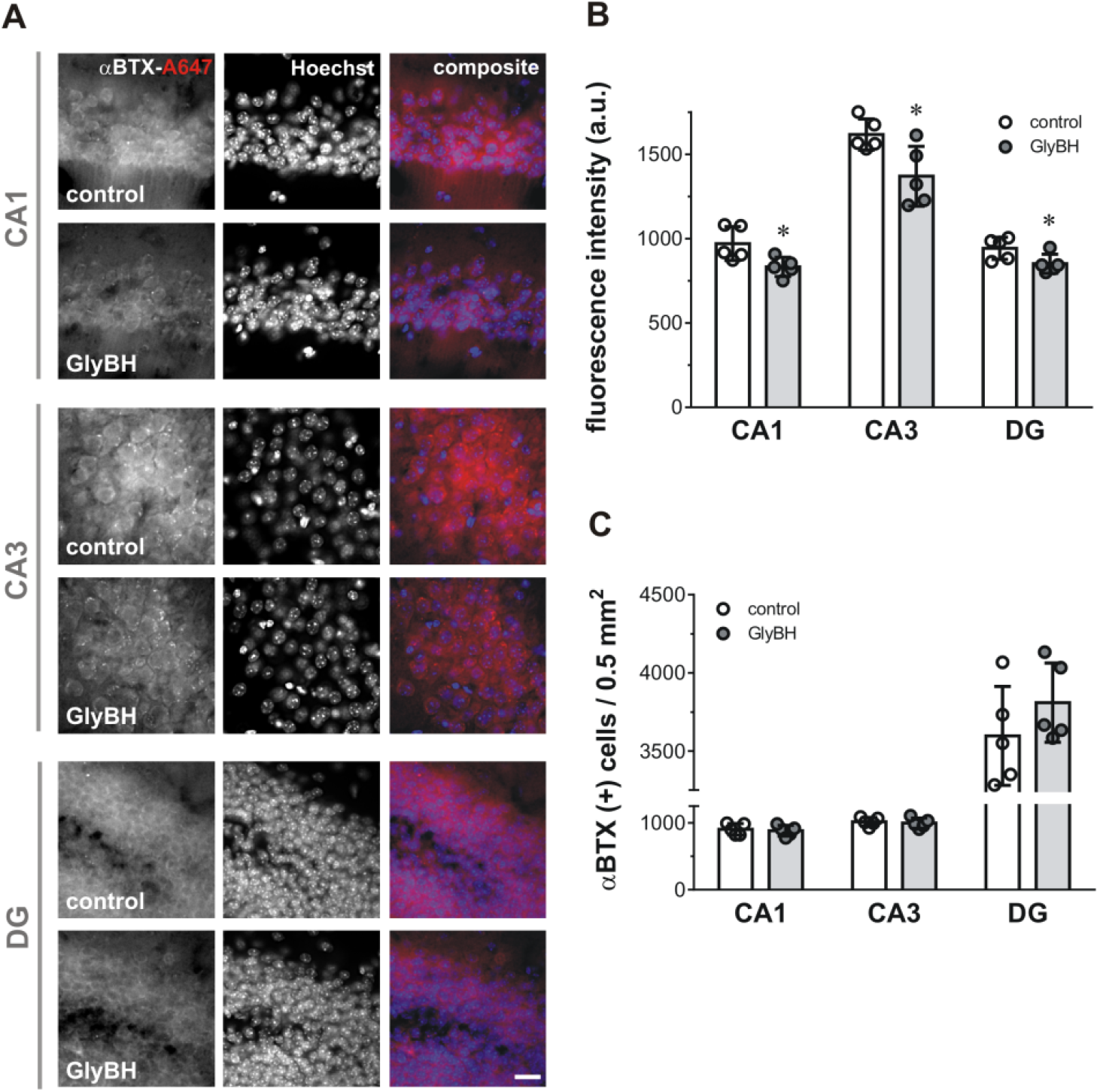
Expression of α7-nAChR in the HPC of control and GlyBH treated mice. **A)** Representative photomicrographs of coronal sections showing αBTX-positive cells in the hippocampal CA1, CA3 and DG regions from control and GlyBH treated mice. α7-nAChR positive cells were labeled using αBTX-A^647^ (left panel). Total cells were visualized by Hoechst stain labelling, (middle panel). A co-localization of both fluorescent channels is shown in the right panel, (*composite*). Bar, 20 μm. **B)**. Fluorescence intensities obtained from αBTX-A^647^ labelling are expressed as arbitrary units (a.u.). **C)** The number of αBTX-A^647^ labelled cells (corresponding to the α7-nAChR expressing cells) was normalized to a fixed area (0.5 mm^2^Bar graphs show mean ± SD, with data points. n = 5 mice per group. * denotes *p* < 0.05, statistically significant different from control mice (Student’s t test).

### 3.6. IN administration of GlyBH did not affect the nigrostriatal pathway

The nigrostriatal pathway plays an essential role in the control of voluntary motor movement (Kuhar et al., 1999). In order to evaluate whether this system was affected by GlyBH treatment or not, we labelled TH-expressing neurons in coronal sections containing the CPu and SN as in Baier et al. (2014). As shown in Fig. 5, there were no differences in enzyme expression in the CPu between control and GlyBH treated mice. Also, the number of TH (+) cells was not affected in any group.

**Figure 5.**
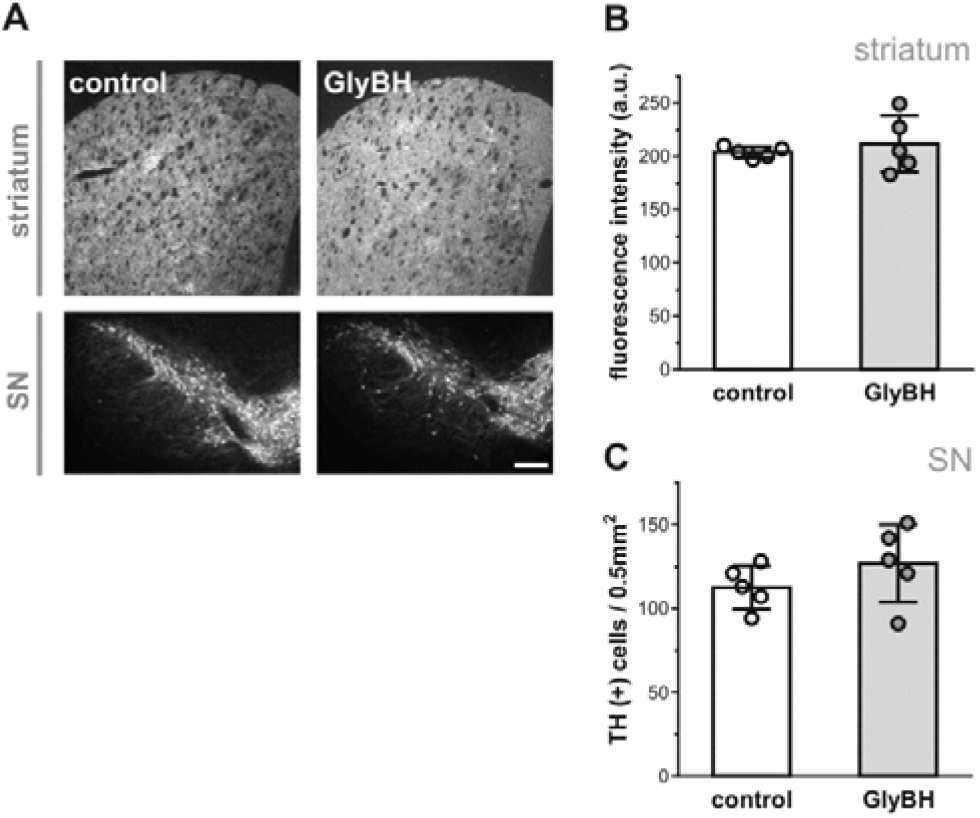
TH expression in the CPu and the SN of control and GlyBH treated mice. **A)** Representative photomicrographs of coronal sections showing TH immunofluorescence in the CPu (upper panel) and in the SN (lower panel) regions from control and GlyBH treated mice. Bar, 250 μm. **B)** Fluorescence intensities from TH labelling were expressed as arbitrary units (a.u.). **C)** The number of TH-expressing cells was normalized to a fixed area (0.5 mm^2^). Bar graphs show mean ± SD, with data points. n = 5 mice per group.

### 3.7. IN administration of GlyBH increased the number of GFAP-immunopositive cells in the anterior olfactory nucleus

Reactive astrogliosis and microgliosis are important features of the neuroinflammatory process (Tristao et al., 2014; Troncoso-Escudero et al., 2018). Quantitative GFAP immunostaining was used to evaluate the number of GFAP (+) cells in different brain areas. As shown in Fig. 6, the region of the AON presented an increase in the FI (p < 0.05) as well as in the number of GFAP (+) cells (p < 0.01) in GlyBH treated mice. No statistically differences were found in the other studied areas (i.e., PFC, CTx, HPC, MS, DBB, CPu and SN, data not shown).

**Figure 6.**
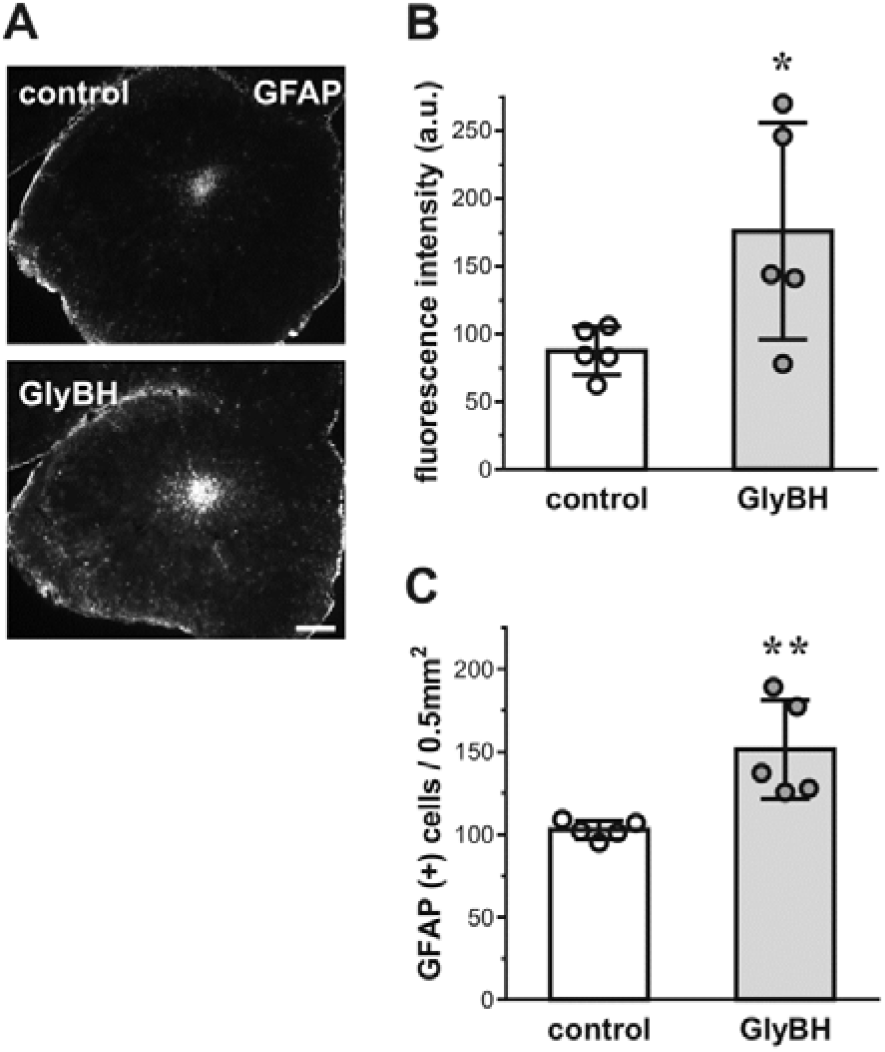
GFAP expression in the AON of control and GlyBH treated mice. **A)** Representative photomicrographs of coronal sections showing GFAP immunofluorescence in the AON region of control and GlyBH treated mice. Bar, 250 μm. **B)** Fluorescence intensities of GFAP sections are expressed as arbitrary units (a.u.). **C)** The number of GFAP-expressing cells was normalized to a fixed area (0.5 mm^2^). Bar graphs show mean ± SD, with data points. n = 5 mice per group.* *p* < 0.05; ** *p* < 0.01 (Student’s t test).

### 3.8. IN GlyBH administration decreased the activity of transaminases in specific brain areas

GPT and GOT enzymes are known regulators of the metabolism of the excitatory neurotransmitter GLU (Daikhin and Yudkoff, 2000; Matthews et al., 2000). In order to investigate whether IN GlyBH administration affects GLU metabolism in the brain, the activity of GPT and GOT was determined in the OB, PFC, CPu, CTx, HPC and Mb. GlyBH treatment resulted in altered activity of the GPT enzyme. Indeed, GPT showed a significant activity reduction in the PFC (p < 0.001) while, in the CPu, its activity was increased (p < 0.05) (Fig. 7A). As for GOT, a significant reduction in enzyme activity was observed in both, the PFC (p < 0.05) and the HPC (p < 0.05) of IN GlyBH administrated mice (Fig. 7B).

**Figure 7.**
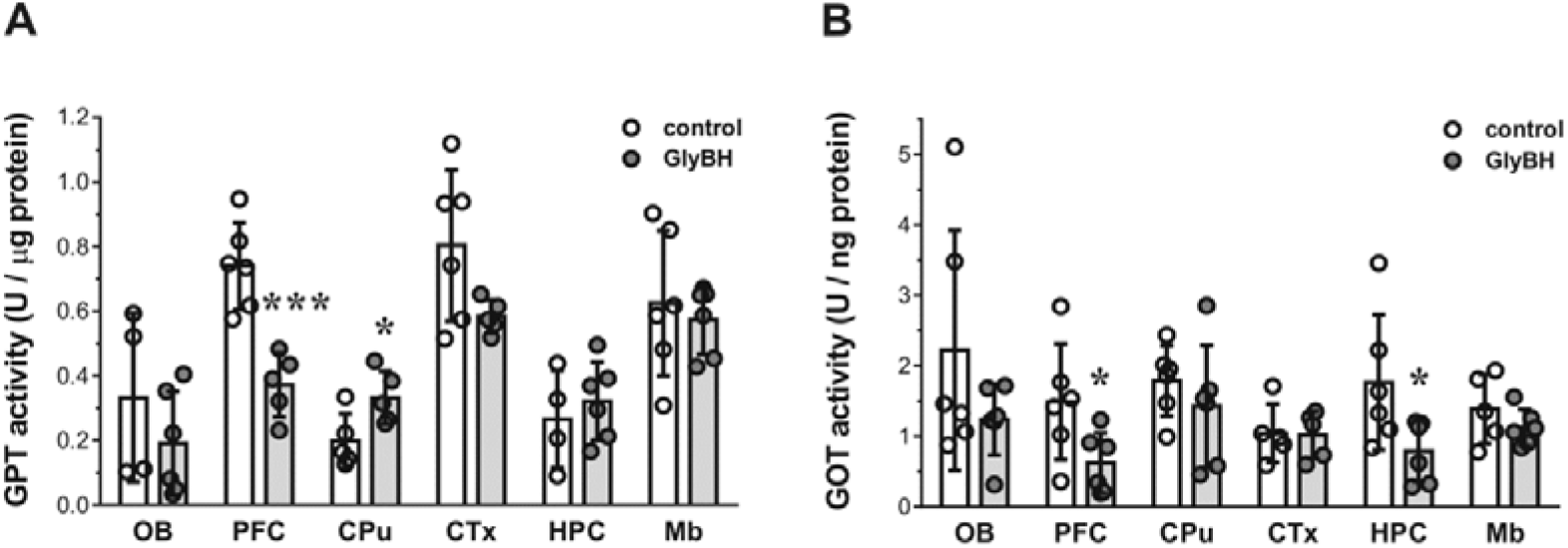
Transaminases activity in different brain areas of control and GlyBH treated mice. **A)** GTP and B) GOT activities in the indicated brain regions from control and GlyBH treated mice. Bar graphs show mean ± SD, with data points. n = 6 per group. * *p* < 0.05; \*\*\**p* < 0.001 (Student’s t test).

## 4. DISCUSSION

The present study revealed that repeated IN administration of a GlyBH in adult mice impairs the redox balance in different brain areas (i.e., OB, PFC, CPu, CTX, HPC and Mb). Furthermore, IN GlyBH exposure modified the activity of different enzymes involved in the cholinergic (OB) and glutamatergic (PFC, CPu and HPC) pathways. Additionally, GlyBH treatment decreased the number of cholinergic neurons in the MS as well as the expression of the α7-nAChR in the HPC. Our results suggest that these alterations could be responsible, at least in part, of the disturbances observed in anxiety, memory and locomotor activity described previously (Baier et al., 2017).

Commercial formulations of pesticides are a mixture of chemicals, composed of an active principle together with inerts or adjuvants, which may have their intrinsic toxicological properties (Mesnage and Antoniou, 2017). GlyBHs are more toxic than Gly alone (El-Shenawy, 2009; Mesnage et al., 2013), and the toxicity of GlyBHs is proportional to their concentration of ethoxylated surfactants (Mesnage et al., 2013). Therefore, it should be noted that the results described in the present study cannot be attributed solely to Gly.

Nerves connecting the nasal passages to the brain and spinal cord, the vasculature, and the cerebrospinal fluid have been implicated in the transport of molecules from the nasal cavity to the CNS (Dhuria et al., 2010; Prediger et al., 2012). Considering that Gly is a glycine analog, this chemical could be a substrate for the glycine-uptake pathways in the OE and thus, reach the CNS through the amino acid transporters LAT1/2 (Xu et al., 2016). Additionally, accompanying adjuvants may contribute to membrane permeabilization, favoring Gly absorption through the OE. Indeed, the addition of adjuvants to GlyBHs enables Gly, a water soluble molecule, to penetrate lipid membranes (Gehin et al., 2005; Mesnage et al., 2019; Williams et al., 2000).

Many xenobiotics that access the brain through the OE could generate free radicals which, in turn, may lead to oxidative stress (Rey et al., 2018), a phenomenon involved in numerous neuropathological processes (Carvalho et al., 2017; Chong et al., 2005; Kim et al., 2015) Exposure to Gly or GlyBH produces oxidative stress in various tissues, including the brain (Beuret et al., 2005; Cattani et al., 2014; El-Shenawy, 2009; Gallegos et al., 2018; Larsen et al., 2012; Modesto and Martinez, 2010). The molecular mechanisms through which Gly and GlyBH induce oxidative stress are well characterized. Uncoupling of mitochondrial oxidative phosphorylation may be a major effect of Gly and GlyBH intoxication (Olorunsogo et al., 1979; Peixoto, 2005; Pereira et al., 2018). The impaired mitochondrial function evoked by GlyBH would be related to increased reactive oxygen species (ROS) production (Bailey et al., 2018; Gomes and Juneau, 2016). Also, Gly or GlyBH exposure result in the alteration of the brain antioxidant system activity (Astiz et al., 2009a, b; Bali et al., 2019; Cattani et al., 2017; Cattani et al., 2014; Gallegos et al., 2018). In proteins, oxidation of free thiol groups may affect its structure, its catalytic activity or its ability to interact with other proteins. Decreased total thiol level was observed in various neurological disorders (McBean et al., 2015; Mungli et al., 2009). IN GlyBH administration decreases total thiol content in the OB, CPu, CTx and HPC. CAT, an essential component of the enzymatic antioxidant defense, is affected in several neurological processes (Niedzielska et al., 2016). Similarly to Gallegos et al. (2018), CAT activity was decreased in the PFC, HPC and Mb of GlyBH-treated mice. In the HPC both, total thiol levels and CAT activity were decreased. These results would indicate that the HPC, which is implicated in the modulation of anxiety and memory, was the brain area most affected by oxidative damage. High levels of free radicals change the expression of nAChRs and N-methyl-D-aspartate receptors, deteriorate cellular membranes by lipid peroxidation, damage receptors by protein oxidation, and modify gene expression by DNA oxidation (Guan, 2008). Increased oxidative stress correlates with impaired memory in the NOR test (Keeney et al., 2018) and with anxiety behaviour in the plus maze test (Bouayed et al., 2009).

Alterations in cholinergic circuitry participate in the cognitive impairments observed in neurodegenerative disorders (Bartus et al., 1982; Maurer and Williams, 2017). In the CNS, cholinergic signal transduction through nAChRs modulates important biological processes such as learning, memory, plasticity, neuroprotection and neurodegeneration (Gotti et al., 2006; Oddo and LaFerla, 2006). The most abundant nAChR subtypes in the brain are the α4β2 and the α7 forms (Gotti and Clementi, 2004; Gotti et al., 2006), which are involved in a wide variety of diseases affecting the nervous system (Baier et al., 2015; Barrantes et al., 2000; Gotti and Clementi, 2004). Cholinergic neurons in the MS, the DBB and the nucleus basalis magnocellularis (NBM) project to the OB, neocortex, HPC, and amygdala (Khakpai et al., 2013). The MS/DBB and NBM cholinergic neurons are implicated in recognition memory (Okada et al., 2015), anxiety and the detection of novelty (Carpenter et al., 2017). Hippocampal injection of the nAChR antagonist mecamylamine produces an anxiogenic action (File et al., 1998), and cholinergic signaling through nAChRs participates in anxiety modulation (Picciotto et al., 2015). Cholinergic neurons from the MS are highly sensible to oxidative stress (McKinney and Jacksonville, 2005). The mitochondrial toxicity induced by GlyBH (Bailey et al., 2018; Gomes and Juneau, 2016; Olorunsogo et al., 1979; Peixoto, 2005; Pereira et al., 2018) could affect the cholinergic neuron physiology, since mitochondrial dysfunction is involved in the beginning of neuronal deterioration (Guo et al., 2017; Lezi and Swerdlow, 2012). The alterations observed in the number of ChAT (+) cells in the MS together with the lower α7-nAChR expression in the HPC could contribute to the anxiogenic behaviour and the recognition memory impairments observed in intranasally GlyBH-treated mice (Baier et al., 2017).

OB is innervated by cholinergic inputs from the basal forebrain, and neurodegenerative disorders related to cholinergic systems are accompanied by olfactory dysfunctions (Attems et al., 2014; Castillo et al., 1999). In a mouse model of Parkinson’s disease the decline of AChE activity in OB precedes that in CTx (Zhang et al., 2015). Interestingly, in the present study IN GlyBH administration decreased the activity of AChE in the OB. Gly is a weak inhibitor of AChE (Larsen et al., 2016) and brain AChE activity was significantly reduced in perinatal GlyBH-exposed rats (Cattani et al., 2017; Gallegos et al., 2018). Chronic inhibition of brain AChE could lead to an excess of acetylcholine (ACh) at the cholinergic synapses, with the consequent excitotoxic damage and degeneration of cholinergic systems (Zaganas et al., 2013). Considering that the time between the last IN GlyBH exposure and AChE activity determination was ~15 days, the AChE activity reduction could be a consequence of several factors including gene and protein expression regulation, enzyme turnover, or even death of AChE competent cells.

Similar to the results obtained after IN paraquat administration (Rojo et al., 2007), IN GlyBH induced decreased locomotor activity without disturbing the nigrostriatal system. Ait Bali et al. (2017) reported that oral chronic exposure to a GlyBH (250 and 500 mg/kg daily for 12 weeks) decreased TH labeling in the SN of treated mice. The difference between the results obtained (besides the different exposure pathway employed) could be attributed to the higher GlyBH doses and exposure time used by Ait Bali et al. (2017).

Reactive astrogliosis and microgliosis are important features of the neuroinflammatory process (Tristao et al., 2014; Troncoso-Escudero et al., 2018). The AON has been proposed as the earliest site of neurodegeneration (Aqrabawi and Kim, 2018; Flores-Cuadrado et al., 2019; Price et al., 1991). Astroglial labeling was higher in the AON of a Parkinson’s disease animal model (Flores-Cuadrado et al., 2019). After IN GlyBH administration the AON showed increased number of GFAP expressing cells. However, no changes in GFAP expression were observed in PFC, CTx, HPC, MS, DBB, CPu and SN in our study. Administration of MPTP increases the GFAP mRNA levels only at early time points (Ciesielska et al., 2009). This acute GFAP response could be responsible for the absence of detectable changes in the number of astrocytes in the other brain areas studied here.

GLU is the main excitatory neurotransmitter in the CNS (Erecinska and Silver, 1990), and plays an important role in spatial learning and memory processes (Dennis et al., 2016; Myhrer, 2003). Once released, GLU must be rapidly removed (Daikhin and Yudkoff, 2000), in order to avoid neurotoxicity (Leibowitz et al., 2012). GLU excitotoxicity is one of the main factors for ROS generation in the brain (Bai et al., 2016; Cattani et al., 2017; Vishnoi et al., 2016). IN exposure to GlyBH affected the GPT activity in PFC and CPu. Besides, the activity of GOT was impaired in PFC and HPC. Perinatal exposure of GlyBH leads to inhibition in the activity of both, GOT and GPT in PFC, CPu and HPC of rat offspring (Gallegos et al., 2018). Cattani et al. (2014) reported inhibition of these transaminases, decrease in glutamine synthase’s activity, along with an increase in GLU release and lower GLU uptake in the HPC of rats exposed to GlyBH. These imbalances in GLU pathways could finally affect hippocampal function.

## 5. CONCLUSIONS

The present study shows that IN exposure to GlyBH induces brain oxidative stress which in turn, could trigger impairment of the septo-hippocampal cholinergic pathway, the increment in astrocyte number in the AON, and the disruption of the activity of enzymes related to the cholinergic and glutamatergic systems in specific brain areas. Then, the disturbances enumerated above could be responsible for the neurobehavioural alterations described previously i.e., decrease in locomotor activity, increase in anxiety state, and impairment in recognition memory. These results strengthen the importance of studying the IN access pathway in relationship to the aetiology of CNS disorders and may help lawmakers to improve life quality, especially for risk population.

## Supporting information

Supplementary Material 1_MSDS Glifoglex

Supplementary Material 2_Table Gly-GlyBH doses

## Acknowledgements

The authors wish to thank Wiener Laboratories for the kind donation of the diagnostic kits and to Dr. Eugenio Aztiria (Instituto de Investigaciones Bioquímicas de Bahía Blanca, Bahía Blanca, Argentina) for his helpful reading of the manuscript.

---

**Supplementary Fig. S1.** W*ater and food intake, and body weight, under control and IN GlyBH conditions.* **A)** Water intake, **B)** food intake, and **C)** body weight at the indicated days of treatment. All data are presented as mean ± SEM. n = 6 for each group.

**Supplementary Material 1.** MSDS of Glifloglex®.

**Supplementary Material 2.** Gly and GlyBH doses used in mice studies.

