## Supplementary Figure S1 for "Intranasal Glyphosate-Based Herbicide Administration Alters the Redox Balance and the Cholinergic System in the Mouse Brain"

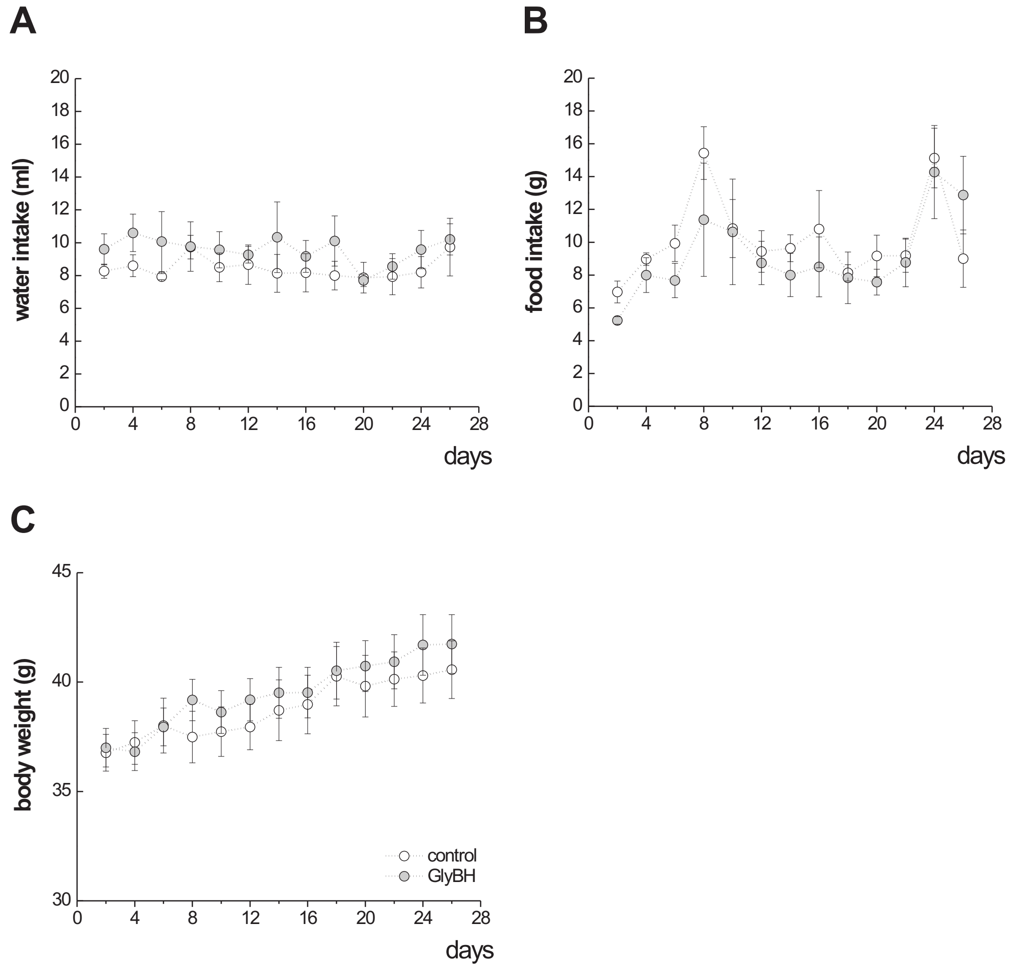


**Supplementary Fig. S1.** W*ater and food intake, and body weight, under control and IN GlyBH conditions.* **A)** Water intake, **B)** food intake, and **C)** body weight at the indicated days of treatment. All data are presented as mean ± SEM. n = 6 for each group.
