## Supplementary Material 1_MSDS Glifoglex for "Intranasal Glyphosate-Based Herbicide Administration Alters the Redox Balance and the Cholinergic System in the Mouse Brain"

### Supplementary Material 1. Material Safety data Sheet (MSDS) Glifoglex®

*English version adapted from Glifoglex® MSDS . Original version, in Spanish, could be found in:*

*https://www.gleba.com.ar/glifoglex?page_id=263* >> *HOJA DE SEGURIDAD*

#### 1. Identification of the chemical product and the company

- **Product identification:** Glyphosate Isopropylamine Salt 48% w/v
- **Recommended uses:** Herbicide
- **Use restrictions:** Use according to the recommendations indicated on the product label.
- **Provider Name:** GLEBA S.A.
- **Supplier Address:** Av. 520 y Ruta Prov. 36 (1903) Melchor Romero, La Plata – Pcia. Buenos Aires - Argentina
- **Provider phone number:** +54-2 214 913 062
- **Emergency telephone number in Argentina:** +54-2 214 913 062
- **Telephone number of Toxicological Information in Argentina:** 0-800-333-0160 - NATIONAL CENTER OF INTOXICATIONS HOSPITAL POSADAS
- **Manufacturer Information:** GLEBA S.A.
- **Web address:** www.gleba.com.ar

#### 2. Hazards identification

- **UN classification:** NU3082-Environmentally hazardous liquid substance, n.o.s. (contains glyphosate isopropylamine salt).
- **Distinctive according to Transport Standard:** 9- Miscellaneous.

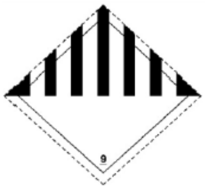


- **Classification according to SGA rev 6:** TOXICOLOGICAL CATEGORY 5 (without pictogram). SERIOUS EYE INJURIES/EYE IRRITATION CATEGORY 2B.
- **GHS label:**
- **Hazard Indications:**
  H303: May be harmful if swallowed.
  H313: May be harmful in contact with skin.
  H333: May be harmful if inhaled.
  H320: Causes eye irritation
- **Precautionary statements:**
  P101: If medical advice is needed, keep the container or label.
  P261: Avoid breathing mists or vapors.
  P264: Wash thoroughly after handling the product.
  P270: Do not eat, drink or smoke while handling this product.
  P271: Use only outdoors or in a well ventilated place.

P273: Do not disperse in the environment.
P304 + P340: IF INHALED: Transport the person outdoors and keep them in a position that facilitates breathing.
P501: Dispose of the contents and/or container in accordance with current national regulations.

#### 3. Composition / information of the components

- **Main components of the mixture:** Glyphosate Isopropylamine Salt
- **Concentration (%):** Glyphosate Isopropylamine Salt 48% w/v
- **Mix Component:**

| **Systematic chemical name** | **Glyphosate Isopropylamine Salt** |
|---|---|
| **Common or generic name** | **2- (phosphonomethylamino) isopropylamine acetate** |
| **Concentration range** | **48% w/v** |
| **CAS number** | **38641-94-0** |

#### 4. First aid

- **In case of inhalation:** Take the patient to a cool and ventilated place. If the person does not breathe grant artificial respiration.
- **In case of skin contact:** Remove clothing and shoes and wash before reuse. Wash the skin with plenty of clean water and thoroughly between hair, nails and skin folds.
- **In case of contact with the eyes:** Wash the eyes with clean or potable water, for at least 15 minutes, taking care that the eyelids are open. In case the affected person uses contact lenses, remove them after the first 5 minutes and then continue with the rinse, in addition the lenses should not be used again.
- **In case of ingestion:** DO NOT induce vomiting. Never give something by mouth to an unconscious person. Take the care center immediately.
- **Expected acute effects:** Nausea, vomiting and diarrhea.
- **Expected delayed effects:** Not described
- **Most important systems / effects:** Not described
- **Protection of those who provide first aid:** Wear gloves
- **Special notes for the attending physician:** No specific antidote is known. Apply symptomatic treatment.

#### 5. Firefighting measures

**Extinguishing Agents:** Use to extinguish flames chemical foam (to avoid ignition of steam) or dry chemical powder.
**Inappropriate extinguishing agents:** Does not apply.
**Products that are formed in combustion and thermal degradation:** Nitrogen oxides, phosphorus oxides, dioxide and carbon monoxide.
**Associated specific hazards:** There is no specific associated danger.

**Specific methods of extinction:** Sprinkle with water to cool off the affected area. Use extinguishing media and signage. Isolate the affected area. The personnel must be using the appropriate foot to fight fire and self-contained breathing apparatus.
**Precautions for emergency personnel and / or firefighters:** The qualified personnel must be affected by a special protective area for firefighting, self-contained breathing apparatus and safety glasses with side shields.

#### 6. Measures to be taken in case of accidental spillage

**Personal precautions:** Do not enter the affected area without adequate protective equipment.
**Protective equipment:** Use equipment detailed in point 8.
**Emergency Procedures:** Isolate the affected sector, people use the appropriate protection elements.
**Environmental precautions:** Contain the spill with inert substances (sand, earth).

**Methods and materials for containment, confinement and / or dejection:** Cover the sewers and prevent accidental spills from reaching water sources. In case of spills on pavements or natural soils contain the spill with inert substances such as vermiculite if available, or sand or dry soil. Subsequently collect the spill in appropriate containers for final disposal. Transfer to an authorized dump for this type of substances, as indicated by the competent authority, another alternative is by controlled incineration in a standard oven at a temperature greater than 1000 ° C with recovery and smoke filtering.

**Cleaning methods and materials:**
- **Recovery:** Recovery does not apply since the substance has been contaminated.
- **Neutralization:** Isolate the affected area, if possible contain the spill with inert substances.
- **Final disposition:** Dispose according to what is indicated by the competent authority.
- **Additional disaster prevention measures:** Prohibit the entry of unauthorized personnel in warehouses, collection or distribution sites.

#### 7. Handling and storage

**Handling**
- **Precautions for safe handling:** Personnel involved in handling the product must use all recommended personal protection elements.
- **Operational and technical measures:** Wash clothes after handling.
- **Precautions:** Do not handle without authorization from the security officer. The product is not flammable, however you should avoid smoking, use of cell phones, lamps and plugs that are not explosion-proof or the use of any element that could generate a spark.
- **Local / general ventilation:** You must have a ventilation system.
- **Contact Prevention:** Wear protective clothing.

**Storage**
- **Conditions for safe storage:** Fresh and dry place, with good ventilation, the products should be stored in shelves separated from the floor. Do not store food and medicine for animal or human use, seeds and any other that comes into direct contact with men and animals.
- **Technical measures:** In authorized deposit and clearly identified containers.

- **Incompatible substances and mixtures:** Incompatible with oxidizing or reducing agents whose pH is greater than 9 or less than 4.
- **Packaging material:** Always keep in the original packaging. Sealed containers, with visible label.

#### 8. Exposure controls / personal protection

**Permissible concentration:**

- **Weighted permissible limit:** Not determined
- **Absolute permissible limit:** Not determined
- **Temporary permissible limit:** Not determined
- **Odorific Threshold:** Not determined
- **Biological standards:** Glyphosate, Aminomethylphosphonic acid
- **Monitoring Procedure:** Blood glyphosate and aminomethylphosphonic acid level.

**Protective elements**

- **Respiratory protection:** Protective mask
- **Hand protection:** Neoprene or latex gloves.
- **Eye protection:** eye protection glasses
- **Skin and body protection:** Full Tyvek suit with hood.

**Engineering measures:**

Handle following all safety measures applicable to the product and the personal protection elements already indicated (8.1.c)

#### 9. Physical and chemical properties

- **Physical state:** Liquid
- **How it is presented:** Soluble concentrate
- **Colour:** Gold yellow
- **Odor:** Not obvious
- **pH:** 4.5 - 5.0
- **Melting point / freezing point:** Not available
- **Boiling point, initial boiling point and boiling range:** Not available
- **Flammability limits (LEL and UEL):** Not flammable
- **Explosive Limit:** Not explosive
- **Vapor pressure:** Not available
- **Vapor Density:** Not applicable
- **Density:** 1,155 g / mL (20 ° C)
- **Solubility (ies):** Soluble in water
- **Partition coefficient n-octanol / water:** Not available
- **Autoignition temperature:** Not available
- **Odor Threshold:** Not obvious
- **Evaporation rate:** Not available
- **Inflammability:** Non-flammable
- **Viscosity:** Not applicable

**10. Stability and reactivity**

- **Chemical stability:** Stable for two years.
- **Dangerous reactions:** Not applicable
- **Conditions to avoid:** Reactive or highly unstable substances.
- **Incompatible materials:** Incompatible with oxidizing or reducing agents whose pH is greater than 9 or less than 4.
- **Hazardous decomposition products:** Nitrogen oxides, phosphorus oxides, dioxide and carbon monoxide

**11. Toxicological information**

**Acute Oral Toxicity:** LD 50 rats> 10,000 mg / kg
**Acute Dermal Toxicity:** LD 50 rats> 10,000 mg / kg
**Acute Inhalation Toxicity:** LC 50 rats: 10.42 mg / L 4 hours
**Irritation / Skin corrosion:** Not irritating
**Serious eye damage / eye irritation:** Mild irritant
**Respiratory or skin sensitization:** Not skin sensitizing
**Cell mutagenicity:** The active ingredient is not mutagenic
**Carcinogenicity:** The active ingredient is not carcinogenic
**Reproductive Toxicity:** The active ingredient is not teratogenic
**Specific target organ toxicity - single exposure:** Not available
**Specific target organ toxicity - repeated exposures:** Not available
**Inhalation Hazard:** Not available
**Related symptoms:** Not available

**12. Ecological information**

- **Ecotoxicity:**
  Birds: > 5,000 mg / kg LD50
  Algae: > 1,000 mg / L 72 hr LC50
  Daphnias: > 600 mg / L 48 hr LC50
  Earthworms: Not available.
  Fish: > 100 mg / L 96 hr LC50
- **Persistence and degradability:** It is mainly degraded by microbiological action. It is unstable in strongly alkaline medium.
- **Bioaccumulative potential:** Low bioaccumulation potential.
- **Mobility in soil:** Low mobility in soils.

**13. Information on final disposition**

- **Waste:** Incineration in Standard type ovens at more than 1,100ºC temperature, 2 ”of residence. Combustion and destruction efficiency: 99.9%
- **Contaminated container and packaging:** Perform triple washing of the containers, render them unusable and send them to the authorized collection center for chipping and subsequent transfer to the dump or recycling. Set the containers in a clearly identified place, until the authority defines the final destination.

- **Contaminated material:** Collect in clearly identified containers, finally transfer to an authorized deposit for this type of substances, for later disposal in accordance with the provisions of the competent authority.

#### 14. Transport information

| | Mode of transport | | |
|---|---|---|---|
| | **LAND** | **MARITIME** | **AIR** |
| Regulations | RID/ADR | IMDG | IATA |
| NU Number | 3082 | 3082 | 3082 |
| Official Designation of Transportation | Dangerous liquid substance for the environment, e.g. (contains glyphosate isopropylamine salt) | Dangerous liquid substance for the environment, e.g. (contains glyphosate isopropylamine salt) | Dangerous liquid substance for the environment, e.g. (contains glyphosate isopropylamine salt) |
| Primary Hazard Classification | 9 | 9 | 9 |
| Secondary Hazard Classification | -- | -- | -- |
| Packing group | III | III | III |
| Environmental hazards | May be harmful to aquatic organisms. | May be harmful to aquatic organisms. | May be harmful to aquatic organisms. |
| Special precautions | Not applicable | Not applicable | Not applicable |

#### 15. Regulatory information

**National Regulations:** IRAM 41400 standard.
**International Regulations:** RID, IATA, IMDG.

**The receiver should pay attention to the possible existence of local regulations.**

#### 16. Other information

**Change control:** Update to SGA rev.6
**Abbreviations and acronyms:**

LD50: Lethal dose 50.
LC50: Lethal concentration 50.
EC: Effective concentration 50.
NOEC: Concentration without observed effect.

**References:** Company Studies
**Validity:** 3 years from the date of update

**It is necessary to have specific training for the handling of the chemical.**
