## Supplementary Material 2_Table Gly-GlyBH doses for "Intranasal Glyphosate-Based Herbicide Administration Alters the Redox Balance and the Cholinergic System in the Mouse Brain"

**Supplementary Material 2:** Gly and GlyBH doses used in mice experiments

| mouse strain | Gly or GlyBH | doses (mg/kg) | days/times | pathway | Author |
|---|---|---|---|---|---|
| ICR | Gly | 400 | 28 d | oral | Gao et al., 2019 [1] |
| Swiss | GlyBH | 250, 500 | 42-84 d | oral | Bali et al., 2019 [2] |
| Kunming | GlyBH | 60, 180, 540 | 35 d | oral | Jiang et al., 2018 [3] |
| Swiss | GlyBH | 250, 500 | 42-84 d | oral | Aitbali et al., 2018 [4] |
| Swiss | GlyBH | 250, 500 | 42-84 d | oral | Aitbali et al., 2017 [5] |
| C57BL/6 | Gly | 200 | single-7d | i.p. injection | Ford et al., 2017 [6] |
| CF1 | GlyBH | 50 | 12 d | IN | Baier et al., 2017 [7] |
| C57BL/6 | Gly and GlyBH | ~ 0.005, 0.05, 5 | 7-9 d | IN (anesthetized) | Kumar et al., 2014 [8] |
| Swiss | GlyBH | 50, 500 | 15 d | oral | Jasper et al., 2012 [9] |
| Swiss | GlyBH | 50 | single | i.p. injection | Cavusoglu et al., 2012 [10] |
| Swiss | GlyBH | 25, 50 | single | i.p. injection | Prasad et al., 2009 [11] |
| Swiss | GlyBH | 25 | 9 d | topical | George et al., 2010 [12] |
| ICR | GlyBH | 600 | single | oral and i.p. injection | Heydens et al., 2008 [13] |
| C57BL | GlyBH | 1080 | single | oral | Dimitrov et al., 2006 [14] |
| Swiss | GlyBH | 50, 100, 200 | 2 d | i.p. injection | Grisolia 2002 [15] |
| Swiss | Gly and GlyBH | 122, 130, 152,182, 270 | 1 d | i.p. injection | Peluso et al., 1998 [16] |
